# Tissues Guide Dependence of Treg on the Transferrin Receptor

**DOI:** 10.64898/2026.02.01.703140

**Authors:** Michelle Montoya, Yasmine T. Toudji, Ata Ur Rehman, Anton Zhelonkin, KayLee K. Steiner, Teresa Tamborra-Walton, Katherine N. Gibson-Corley, Samantha St. Jean, Denis A. Mogilenko, Jeffrey C. Rathmell, Kelsey Voss

## Abstract

Activated T cells increase transferrin-bound iron uptake via the transferrin receptor, also called CD71. We previously demonstrated that targeting CD71 with an antibody to reduce iron update can modify CD4 T cell function, with different effects on T_H_1, T_H_17, and regulatory T (Treg) cells. CD71 blocking antibody-treated Tregs had no loss of viability or differentiation, and Foxp3 expression was increased. However, a genetic deletion of *Tfrc* (the gene for CD71) driven by *Foxp3*-Cre was reported to cause a lethal autoimmunity. Whether altered immune homeostasis or insufficient early developmental tolerance drive the phenotype of CD71 knockout (KO) Treg mice were unclear. Here, we examined the *Foxp3*-^YFP^-Cre KO mouse model and a tamoxifen-inducible KO model in adults to determine the role of CD71 expression in Treg cells. We hypothesized that due to a lack of iron for mitochondrial metabolism, KO Treg adapt to rely heavily on glycolysis and become unstable, promoting pro-inflammatory exTreg cells. This effect was not universal, however, and necropsy analyses revealed tissue-specific inflammation. While the colons of mice with KO Treg cells appeared healthy, skin and lung tissue were severely inflamed. Metabolically, KO Treg cells had a significant decrease in their glycolytic capacity and instead increased oxidation of amino acids and fatty acids. In inflamed skin, which that promotes increased oxidative stress, CD71 expression in Treg cells suppressed tissue inflammation in a model of atopic dermatitis-like disease. These results indicate the CD71-iron axis as a new immunometabolic regulator of Treg cell functions in immune and non-immune organs.

**Capsule Summary:** A loss of the transferrin receptor in Tregs causes severe autoimmunity and here we clarify how Tregs rely on this receptor for iron in specific tissues and disease settings including atopic dermatitis.

## Introduction

T lymphocytes (T cells) depend on metabolic processes including iron metabolism for proliferation and effector functions (1–3). This metabolism is also integral in determining the differentiation of various CD4 T helper (T_H_) cell subsets (4), and consequently, targeting metabolism offers a powerful tool to modulate immune responses. During activation, T cells increase transferrin-bound iron uptake via the transferrin receptor, also called CD71. This iron uptake is important for proper T cell activation, as loss-of-function mutations in the gene for CD71 (*TFRC* in humans, *Tfrc* in mice) results in combined immunodeficiency (CID) with poor T cell proliferation (5, 6). Along with insufficient iron, excessive iron can also be damaging, disrupting redox balance and increasing reactive oxygen species (ROS) and lipid peroxides that promote ferroptosis (7, 8).

We previously demonstrated that targeting CD71 with an antibody which lowers cellular iron can modify T cell function, with different effects on T_H_1, T_H_17, and regulatory T cells (Treg) (9). Of note, Treg cells treated with anti-CD71 had no loss of viability or differentiation, and the master transcription factor Fork Box P3 (Foxp3) expression was increased. However, a genetic deletion of *Tfrc* in Treg cells driven by *Foxp3*-Cre was reported to result in a lethal autoimmune phenotype in mice (10, 11). These findings highlight a sensitive balance between iron levels and immune cell function and suggest that optimal levels of CD71 expression and iron in Treg cells are essential for immune homeostasis.

Despite the severe phenotype of CD71 knockout (KO) Treg cells in mice, the mechanisms responsible for maintaining immune homeostasis in this model have been difficult to study. Although KO Treg cells underwent normal development in the thymus, they failed to effectively proliferate peripherally during early development (10). However, whether defective proliferation is solely responsible for their failure to control inflammatory T cells is unclear. Interestingly, there was a higher reliance of tissue Treg cells on CD71 expression compared to those in lymphoid organs. Certainly, different tissue microenvironments and iron availability at different locations could influence these phenotypes. Because Treg cells regulate autoimmunity and tissue homeostasis across varying tissue types, we sought to explore in which microenvironments Treg cells rely on CD71-mediated iron metabolism for their function.

The role of CD71 expression in Treg cells in establishing tolerance early in life versus steady-state immune homeostasis in adulthood also requires further clarification. Here, we examined both the *Foxp3*-^YFP^-Cre knockout (KO) mouse model as well as a tamoxifen-inducible model in healthy adult mice to determine the role of CD71 expression in Treg cells within this context. Additionally, Treg metabolism is inherently tied to functional stability and Foxp3 expression (12–14). Therefore, we investigated how KO Treg cells metabolically adapt to compensate for a loss of transferrin-bound iron. We hypothesized that due to a lack of iron that supports mitochondrial metabolism, KO Treg cells become unstable due to metabolic adaptations that rely heavily on glycolysis, destabilizing their Foxp3 expression and promoting “ex” Treg cell differentiation (15). We also assessed if iron-dependent phenotypes of Treg cells are exacerbated in tissue microenvironments that promote increased oxidative stress or have low iron availability.

## Methods

### Mice

All animal procedures were approved by the Institutional Animal Care and Use Committee (IACUC) of the University of Virginia and/or Vanderbilt University, conducted in accordance with humane guidelines. Male and female mice aged 8-12 weeks were housed under standard conditions with ad libitum access to food and water. Mice were monitored daily for signs of distress, weight loss, or adverse reactions. Animals were euthanized using CO₂ inhalation followed by cervical dislocation in accordance with AVMA guidelines.

CD4-Cre;*Tfrc* floxed mice were previously described (9). CD4-Cre negative mice homozygous for floxed *Tfrc* were then used to breed to either Foxp3^YFP^-Cre mice (Jax stock # 016959) or Foxp3^eGFP^-Cre-ER^T2^ mice (Jax stock # 016961). Heterozygous females positive for Foxp3^YFP^-Cre were considered WT mice. Males heterozygous for *Tfrc* floxed alleles were also considered WT mice. Foxp3^eGFP^-Cre-ER^T2^ mice were all bred to homozygosity for both *Tfrc* and Foxp3^eGFP^-Cre-ER^T2.^ WT and KO status was determined by the injection regimen which was mixed within the same cages.

### Disease models

Colitis model: 6 week-old *Rag1-/-* mice were purchased from the Jackson Laboratory (stock #002216) and acclimated for 2 weeks. WT and KO Tregs were isolated from CD4-cre + and – spleens according to the (Stem Cell Technologies) and mixed in a 1 to 3 ratio with naïve CD4 T cells isolated from WT CD45.1 mice according to the Naïve CD4 T Cell Isolation Kit (Miltenyi Biotec). A total of 1 million cells was transferred into each recipient mouse by IP injection. Monitoring of disease progression was performed by following body weights post-injection and tissue scoring of colon pathology as previously described (16).

Tamoxifen-inducible knockout model: Foxp3^eGFP^-Cre-ER^T2^ mice: Tamoxifen (Sigma-Aldrich, 10540-29-1) was prepared in corn oil at a concentration of 20 mg/mL by rotating at room temperature until fully dissolved. The tamoxifen solution was protected from light, aliquoted and stored at −20°C. Mice were injected intraperitoneally (IP) with 100uL tamoxifen or corn oil vehicle over the course of 2 weeks, 5 injections total. All injections were performed using sterile techniques with a 1 mL syringe and 27-gauge needle.

MC903-induced experimental atopic dermatitis: Homozygous Foxp3^eGFP^-Cre-ER^T2^ /Tfrc^flox/flox^ littermates aged 8-12 weeks were treated with tamoxifen or mock for two weeks followed by treatments of the skin at inner part of ears with ethanol EtOH (10 ul) or Calcipotriol (MC903, 0.1 mM in 10 ul EtOH, Cat#2700 TOCRIS Biotechne) every day for 10 days. On day 10, we profiled itching behavior in all MC903-treated mice in individual cages. The observations were recorded for one mouse at a time for a period of 10 min after 1 min of adjustment period to the new cage. Mice were euthanized at day 11 of the AD model with CO_2_ asphyxiation followed by cervical dislocation. We recorded body weights, performed cardiac puncture to collect blood, and dissected the ears and collected parotid lymph nodes. Ears were photographed, and AD pathology scores were recorded by a blinded dermatologist. Left ears and respective parotid lymph nodes were fixed in 4% PFA and embedded in the OCT cryomedium for hematoxylin and eosin (H&E) histological analysis. Right ears and respective parotid lymph nodes were digested in a collagenase D solution and used for immune cell isolation and phenotyping.

### RNA sequencing

Lungs and spleens were dissected from WT and KO Foxp3-YFP Cre mice after cardiac prefusion with cold PBS. Lungs were diced and digested in DMEM + 2% serum + 0.2% collagenase D for 45 minutes with shaking at 37°C. Cell pellets were strained through 70um filters, lysed with ACK lysis buffer, and resuspended in phenol red-free RPMI media for cell sorting by FACS. Samples were gated on lymphocytes, live, single cells, CD4+, and then either positive or negative for Foxp3-YFP for collection. Treg (YFP+) and Tcon (YFP-) samples were then cryopreserved in heat-inactivated serum + 10% DMSO or lysed immediately for RNA extraction using the Qiagen RNeasy Micro Kit according to the manufacturer’s instructions. Input RNA was subjected to the Nugen Ovation kit to amplify the RNA prior to cDNA synthesis.

Paired-end FASTQ files for each sample were aligned to mouse genome GRCm39 RefSeq assembly GCF_000001635.27 (Release Feb, 2024). We used STAR(17) in two-pass mode (STAR --twopassMode Basic) to align the reads to the reference genome and sort them by coordinate. Each sample was evaluated according to a variety of both pre- and post-alignment quality control measures with FastQC 0.12.1, Picard (Broad Institute) 3.2.0, RSeQC (18) 5.0.4, MultiQC (19) 1.25. Reads were trimmed with Trimmomatic (20) 0.39. PCR duplicates were marked with Picard tools, and the deduplication effect was tested downstream. Aligned read counts were calculated by featureCounts(21) from the subread 2.0.2 release. Bulk RNA-seq count data were analyzed in R 4.4.2. Differential expression analysis was performed using edgeR (22, 23) 4.2.2 and limma-voom (24) 3.62.2. Count data were filtered for lowly-expressed genes using filterByExpr() and library sizes were normalized using the Trimmed Mean of M-values (TMM) method.

### RNA-sequencing batch correction and model fitting

Analysis consisted of both cryopreserved and fresh Treg samples. Therefore, we implemented a two-model comparison approach to assess the impact of processing method (fresh vs cryopreserved) on differential expression results. The first model (Model 1) was cryopreserved-only analysis. To eliminate processing-related confounding, we restricted the analysis to cryopreserved Treg samples and fit a no-intercept interaction model of genotype and organ using limma-voom : *y_g_=X β_g_+ε_g_, X=[ Lung_KO, Lung_WT, Spleen_KO, Spleen_WT]* where each indicator column equals 1 for samples in that organ×genotype group (0 otherwise; no intercept). This model compared KO vs WT within each organ without confounding from processing. The second model (model 2) was a batch-corrected full dataset. To leverage all samples while accounting for processing, we fit a no-intercept voom–limma model with four organ×genotype indicators and one sample-level processing covariate (with fresh coded 1 for fresh and 0 for cryopreserved): *y_g_=X β_g_+ε_g_ , X=[ Lung_KO, Lung_WT, Spleen_KO, Spleen_WT, fresh]*.

Linear model fits were performed using edgeR’s *voomLmFit()* without sample weights. We initially tested empirical sample quality weight estimation *voomLmFit(…, sample.weights = TRUE)* to account for increased variability in the KO group, but this approach downweighted KO samples showing largest deviations from WT controls. While sample weighting yielded more statistically significant genes, it systematically reduced effect sizes toward zero, suggesting the algorithm was treating biologically relevant differences as technical outliers rather than genuine signal. We therefore proceeded without sample weights to preserve the biological signal of interest. Empirical Bayes moderation was applied using *eBayes()* with *robust=TRUE* for improved variance estimation. Key findings from model comparisons: Fold-change agreement: Bland-Altman analysis revealed good agreement for genes with moderate effect sizes (*|mean log-FC| < 2*), but Model 1 systematically inflated fold-changes for strongly dysregulated genes (*|mean log-FC| > 5*), suggesting over-estimation when batch effects are ignored. Variance stabilization: Mean-variance plots showed Model 1 exhibited extreme variance inflation at high expression levels, with several highly expressed genes showing disproportionate variability. Model 2 effectively stabilized these variance estimates by accounting for processing-related technical variation. Statistical power: P-value distributions revealed that Model 1 suffered from reduced statistical power, with a substantial proportion of genes becoming untestable (p-value = 1) due to zero estimated variance in the smaller cryopreserved-only subset. Model 2 rescued these genes by leveraging additional samples. Differential expression robustness: While Model 1 identified more differentially expressed genes overall, only approximately one-third of these calls remained significant in Model 2, suggesting that many Model 1 hits reflected inflated estimates due to reduced sample size rather than genuine biological effects. Importantly, no genes changed direction between models, indicating that batch correction attenuated rather than inverted biological signals. Based on these diagnostics, we selected Model 2 for final analysis as it provided more robust effect size estimates, better variance stabilization, and improved statistical power by leveraging all available samples with appropriate batch correction.

### Gene set enrichment analysis

Gene set enrichment analysis (GSEA) (25) was performed using clusterProfiler(25–27) 4.14.6 with fgsea(28) as the calculation method. All genes passing the low-expression filter ranked by moderated t-statistic were included in the gene list to avoid bias in pathway analysis (29, 30). Genes were ranked by moderated t-statistics from the differential expression analysis. We tested against multiple pathway databases from MSigDB (30) via msigdbr 10.0.2 including: Hallmark gene sets (30) (H), Gene Ontology Biological Processes (C5:GO:BP) (31), Gene Ontology Molecular Functions (C5:GO:MF), KEGG pathways (32) (C2:CP:KEGG), Reactome pathways (33) (C2:CP:REACTOME), WikiPathways (C2:CP:WIKIPATHWAYS), BioCarta pathways (C2:CP:BIOCARTA). GSEA was performed with 100,000 permutations for robust p-value estimation. Pathways with FDR-adjusted p-value < 0.05 were considered significantly enriched. In the dotplots the pathway bubbles outlined with a black outline are those crossing the significance threshold. All processed data, analysis scripts, and bioinformatics analysis results are available at GitHub https://github.com/MogilenkoLabVUMC/KelseyVoss_Tregs/tree/v1.0.0 repository.

### Flow cytometry

Tissues were digested into single cell suspensions by the following methods: Spleens were smushed through a 70-micron filter (Miltenyi Biotec), washed with complete RPMI1640 media (10% heat-inactivated (HI) serum + 1% anti-anti) and then subjected to ACK lysis for 1 minute. Cells were resuspended in FACS buffer (PBS + 2% HI serum + 2mM EDTA) and subjected to various downstream staining procedures. Ear samples were placed into DMEM + 2% HI FBS + 3ug/mL collagenase D and minced into ∼1mm^2^ pieces. Minced samples were digested for 1 hour at 37°C with shaking, and washed through a 70-micron filter with FACS buffer. Pellets were resuspended in PBS + 30% Percoll and subjected to density centrifugation on top of PBS + 80% Percoll for 30 minutes at 4°C, 1500rpm with no brake. The interphase layer was transferred to a new tube, washed with FACS buffer and used for downstream staining protocols. For lung samples, mice were first perfused with cold PBS through the right and left ventricles of the heart. Lungs were then dissected and placed into DMEM + 2% HI FBS + 0.2% collagenase D solution and minced into ∼1mm^2^ pieces. Samples were digested at 37°C with shaking for 45 minutes, passed through a 70-micron filter and washed with FACS buffer. Pellets were ACK lysed for 3 minutes at room temperature, washed with FACS buffer again and then used for downstream staining procedures.

For cytokine staining, cells were resuspended in media instead of FACS buffer, and then restimulated for 4 hours in complete RPMI media + GolgiPlug and GolgiStop (BD Biosciences) + 0.5ug/mL ionomycin + 0.25ug/mL PMA. In all cases, cells were washed with ice cold PBS and incubated with TruStainFcx PLUS solution (Biolegend #156604) for 10 minutes on ice. Surface staining was performed with 50uL of a fluorochrome-conjugated antibodies mastermix specific for targeting immune cell surface markers. Cells were set to incubate in ice for 30 minutes, followed by a wash with FACS buffer. For transcription factor staining, cells were then fixed and permeabilized using the Foxp3 Transcription Factor Staining buffer kit (TNB-0607) and incubated overnight with intracellular antibodies. For cytokine staining, cells were fixed for 30 minutes with Cytofix/Cytoperm buffer (BD Cat # 554722), washed with FACS buffer, and then intracellular staining performed with BD Perm/Wash buffer (BD Cat #554723).

Single-cell energetic metabolism by profiling translation inhibition (SCENITH):

Cell viability of lymph node cell samples was measured by counting the cell suspension using a trypan blue exclusion assay (Bio-Rad TC-20) and resuspended in mouse plasma-like medium (VIMPCS) at a concentration of 1.0×10^6 cells/mL. Cell suspensions were plated into a 96-well U bottom plate. Consequently, the plate was centrifuged, and the supernatant was aspirated. Cells were then resuspended using 180uL of warm VIMPCS media and incubated for 15 minutes at 37°C prior to the addition of inhibitor treatments. Following the initial incubation, cells were subjected to one of four separate inhibitory conditions (10μL) to selectively block metabolic pathways: DMSO (vehicle control), 2-Deoxy-D-glucose (2-DG; 100mM, glycolysis inhibitor), oligomycin (1.5uM, oxidative phosphorylation inhibitor), or a combination of 2-DG and oligomycin (2-DG+O). Each inhibitor was added individually or in combination (2-DG +O), and cells were incubated for 20 minutes at 37°C. Subsequently, 10uM puromycin was introduced to all wells (excluding the non-puromycin controls) and incubation continued for an additional 30 minutes. To maintain volume and nutrient consistency across wells, an additional 10uL of VIMPCS media was added to non-puromycin controls, puromycin controls, and wells only receiving one inhibitor. Samples were analyzed using a 4-laser Cytek Aurora flow cytometer, collecting at least 50,000 events of live cells per condition.

### Pathology

Mice were euthanized by carbon dioxide and secondary cervical dislocation. Complete necropsies were performed by harvesting the brain, cecum, colon, ear (external and middle), femur, heart, kidney, liver, lungs, lymph nodes, pancreas, salivary glands, skeletal muscle, small intestine, spleen, stomach, thymus, and urinary bladder. Tissues were fixed in 10% neutral-buffered formalin, processed in alcohol and xylene, embedded in paraffin, sectioned to a thickness of 5 μm, and stained with hematoxylin and eosin (H&E). The femur was decalcified prior to processing.

### Data Availability

All processed data, analysis scripts, and bioinformatics analysis results are available at GitHub: https://github.com/MogilenkoLabVUMC/KelseyVoss_Tregs/tree/v1.0.0 repository.

## Results

### Tfrc KO Treg cells maintain their suppressive capacity in the gut but not skin

Due to the limited number of KO Treg cells, testing their intrinsic suppressive capacity has been difficult. In one study, KO Treg cells were transferred into *Rag1*^-/-^ mice in a model of intestinal inflammation to test their suppressive function, and KO Treg cells were unable to prevent weight loss and tissue damage compared to wild-type (WT) Treg cells (11). However, the environment in which Treg cells are isolated from for these experiments could influence their phenotype independently of *Tfrc* deletion. To address this, we took advantage of the *CD4*-Cre *Tfrc*^fl/fl^ mouse model, which produces KO Treg cells but shows no signs of inflammation, likely due to the inability of conventional T cells (T_conv_) to proliferate (**Supplemental Figure 1**). To test KO Treg cell suppression, CD25^+^ Treg cells were isolated from Cre^+^ or Cre^-^ mice and transferred into the *Rag1*^-/-^ model of colitis (**Figure 1A**). Although mice that received KO Treg cells started to lose some body weight near the end of the disease course (**Figure 1B**), the pathology in WT versus KO Treg recipient colons was not apparent (**Figure 1C**). Analysis by a board certified veterinary anatomic pathologist on the proximal, middle, and distal regions of the colons revealed significant suppression of disease pathology by WT Treg cells compared to the no Treg control recipients (**Figure 1D-F**). Surprisingly, KO Treg cells also significantly reduced disease, although not to the same extent as WT Treg cells. Together, these data suggest that despite a defect in proliferation, KO Treg cells maintain their suppressive capacity in a colitis model.

**Figure 1.**
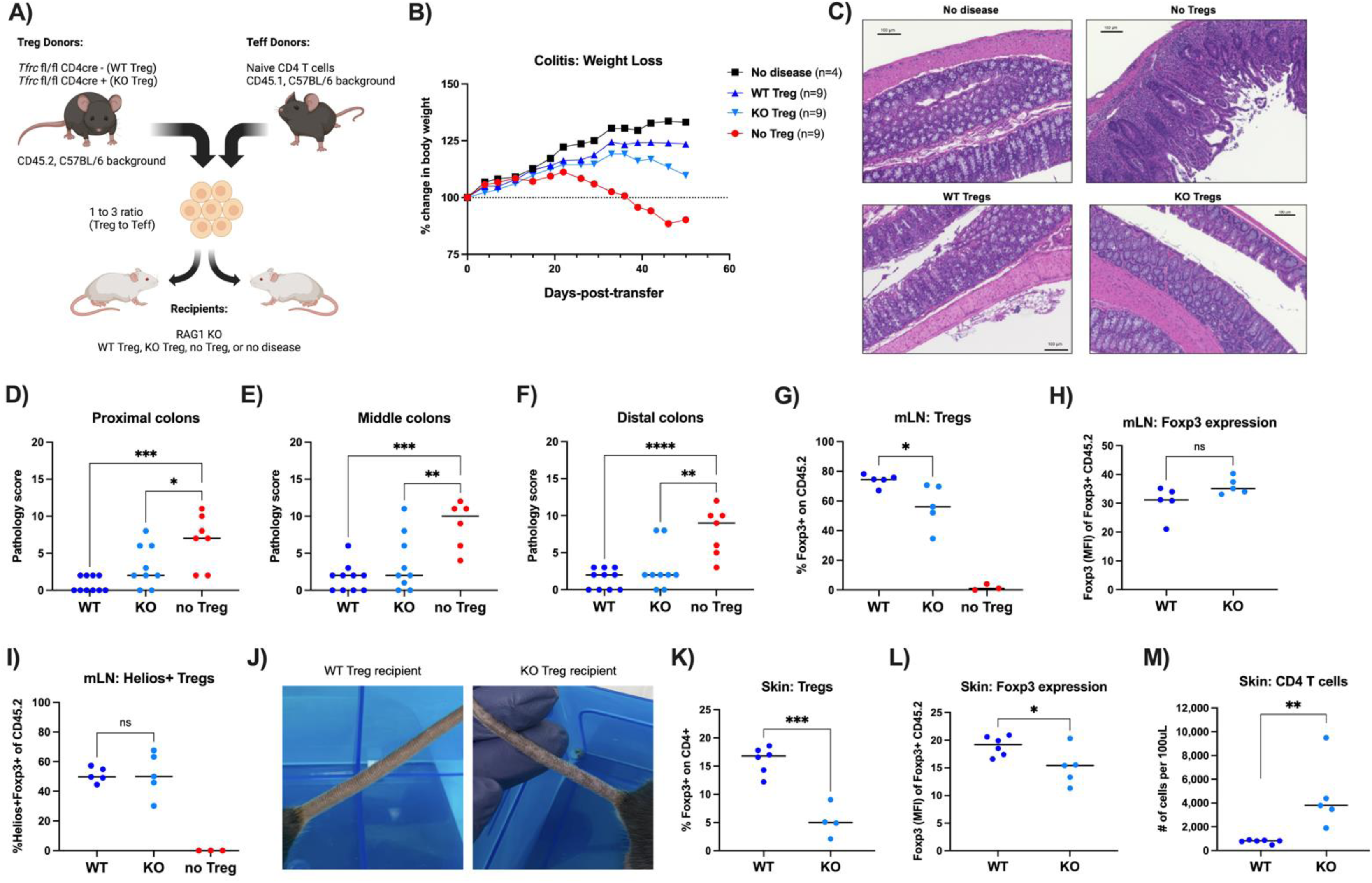
*Tfrc* in Tregs is not required to suppress colitis but is required to prevent skin inflammation. A) To test Treg suppressive function *in vivo*, WT and KO Tregs isolated from CD4-Cre *Tfrc* floxed mice were directly compared in a colitis model. B) Body weight of recipient mice was monitored during disease development until the humane end point when the first mouse had lost more than 20% of its original body weight. Average weight loss of all mice in each group is shown. C) Colons were processed for H&E staining and pathology scores were determined by a blinded veterinary pathologist for the proximal (D), middle (E), and distal (F) regions of the colon. Data are from 2 independent experiments. One-way ANOVA with Sidak’s multiple comparisons test. G) Mesenteric lymph nodes (mLN) were processed for flow cytometry to analyze the frequencies of transferred Tregs, their Foxp3 expression (H), and Helios expression (I). J) One representative tail from both recipient mice of the WT and KO Tregs. K) Ears of recipient mice were processed for flow cytometry to count Tregs (K), Foxp3 expression (L), and quantify CD4 T cells in the skin (M). (K-M) Unpaired two-tailed Student’s t-test.

We next investigated the gut-draining mesenteric lymph nodes (mLN) for Treg homing and stability. In WT Treg recipients, around 80% of the transferred cells identified by CD45.2 retained their Foxp3 expression in the mLN at the study endpoint (**Figure 1G**). However, some KO Treg cells had a significant loss of Foxp3 expression, ranging between 40 and 70%. The intensity of Foxp3 expression, however, was not different between WT and KO Treg cells (**Figure 1H**). Additionally, the percentage of Helios^+^ Treg cells within Foxp3^+^ Treg was not affected in KO Treg, suggesting that Treg cells that resided in the mLN retained some markers of stability during colitis (**Figure 1I**).

Despite similar disease suppression in the colon, we noticed that recipients of KO Treg cells developed gross visual inflammation of their skin, while mice that received WT Treg cells did not (**Figure 1J**). This suggested that *Tfrc* expression may be differentially required for Treg cells in different tissue niches. Skin samples were digested for flow cytometric analyses, revealing that KO Treg recipients had significantly fewer Treg cells in the skin (**Figure 1K**) and those cells which were Foxp3^+^ had less intensity of Foxp3 expression (**Figure 1L**). Indeed, skin samples from KO Treg recipients had increased CD4^+^ T cell inflammation compared to WT Treg recipients (**Figure 1M**). Together, these results suggest that KO Treg cells can suppress inflammation in the gut but not the skin of *Rag1^-/-^* mice.

In line with this, we compared CD71 expression on WT transferred Treg cells among different tissue sites and saw a range of CD71 expression with skin Treg cells having the least CD71 expression compared to splenic Treg and mLN Treg cells having no significant difference (**Supplemental Fig 2A**). Indeed, re-analysis of previously published single-cell RNA sequencing data from tissue Treg showed high levels of *Tfrc* expression in skin-derived Treg cells (**Supplemental Fig 2B**), likely due to a compensatory increase in transcript stability due to low iron availability in skin which regulates *Tfrc* mRNA through RNA-binding proteins (34, 35). Together, these data support the notion that tissue-specific niches with different transferrin-bound iron availability may dictate KO Treg cell stability and function.

### Tfrc expression in Treg cells is required in different tissues

To further investigate the role of Treg-specific *Tfrc* expression in different tissues, we generated *Foxp3*-^YFP^-Cre KO mice. The *Foxp3*-Cre mouse has been generated before and is lethal (11). Our phenotype was similar in many ways, including a failure to thrive with death by around 3.3 weeks of age. This phenotype is characterized by a reduced body condition with body mass and size of the Cre^+^ mice being nearly half that of their WT littermates **(Supplemental Fig. 3A and B)**. Further analyses revealed significant splenomegaly in KO mice (**Supplemental Fig. 3C**). Flow cytometry confirmed prominent T cell lymphoproliferation **(Supplemental Fig. D and E)** with a concomitant reduction in undifferentiated, naive CD4^+^ T cells in KO mice **(Supplemental Fig. F).**

Consistent with previous findings, KO mice revealed a dramatic reduction in Foxp3^+^ Treg cells, although Treg cells were detectable at around 3 to 5% of the splenic CD4^+^ T cells (**Supplemental Fig. 3G**). Within the B cell compartment, KO mice exhibited a large reduction of splenic B cell frequencies **(Supplemental Fig. 3H)**; yet, a larger percentage of these B cells differentiated into plasma cells **(Supplemental Fig. 3I),** which could support an autoimmune-like phenotype (36). In line with this, we also noted a significant increase in the frequency of T follicular helper (T_FH_) cells in the KO mice **(Supplemental Fig. 3J)** which are essential for B cell activation and the manifestation of autoimmunity.

To our surprise and in contrast to previous reports (11), our KO mice showed no signs of gut pathology; a board certified veterinary anatomic pathologist analyzed 10 colon samples from 3-week-old mice (5 WT and 5 KO), scoring them all with a pathology score of “0”. Instead, the most notable inflammatory pathology in our KO mice occurred in the skin and lungs. Around 2 weeks of age, our KO mouse began to show signs of dry, inflamed skin around the base of the tail, which rapidly spread throughout, eventually causing the appendage to become stiff and frail (**Figure 2A**). A similar pattern was observed for the paws and ears, where the KO mice exhibited irritated, flaky patches of skin (**Figure 2A**). Microscopic analysis of the external, middle, and inner ears revealed multifocal bilateral subepithelial otitis externa of the deep external ear canal, composed of primarily neutrophils with fewer macrophages, eosinophils, and lymphocytes (**Figure 2B)**.

**Figure 2.**
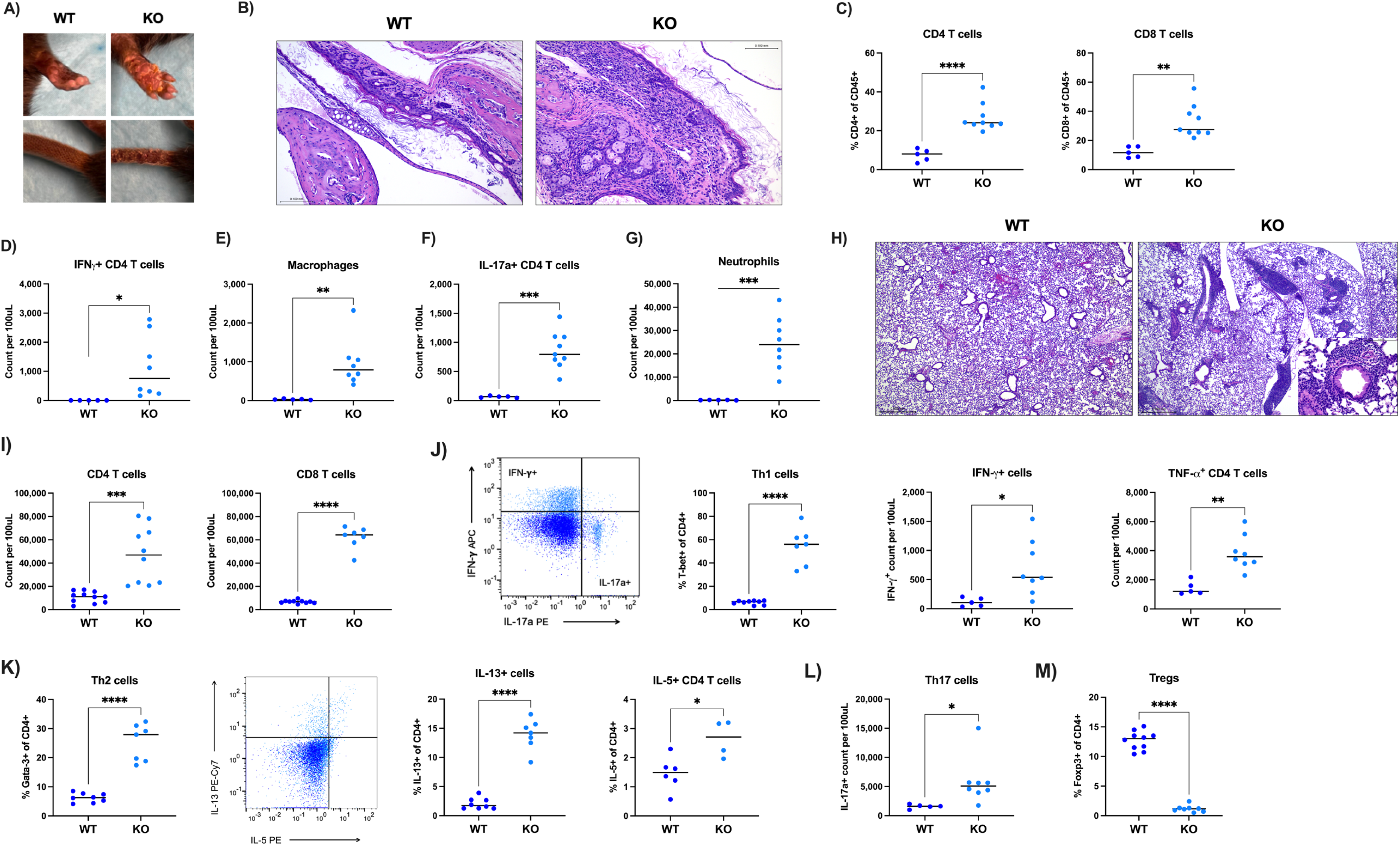
Tissue-independent inflammation of the skin and lung in KO mice. A) Representative images of Foxp3^YFP^-Cre, *Tfrc* floxed mice with scaly skin on the tail and paws of KO mice. B) Representative images of H&E-stained ears from littermates. Scale bar= 0.10 mm. C) Ears were processed into single cell suspensions and CD4+ and cytotoxic CD8+ T cells quantified by flow cytometry. D) IFNy+ CD4 T cells quantified by flow cytometry in the skin samples. E-G) Increased infiltration of macrophages (CD11b+F4/80+), IL-17a⁺ CD4 T cells, and neutrophils (CD11b+Ly6G+) in KO ears. H) H&E-stained lung sections. Scale bar = 0.5 mm, inset scale bar= 0.1 mm. (I) Lungs were processed into single cell suspension to quantify CD4 and CD8 T cells by flow cytometry. J) Frequency of T-bet+ Th1 cells in lungs, IFN-y producing T cells and TNF-a+ T cells. K) Th2-associated populations, Gata-3+ Th2 cells, IL-13 and IL-5 cells. Frequency of Th17 cells (L) and Foxp3+ Tregs (M) in the lungs. All statistical tests are unpaired two-tailed Student’s t-test.

Flow cytometric analyses of cells isolated from ears showed elevated T cell counts of both CD4^+^ and cytotoxic CD8^+^ T cells (**Figure 2C)**. Specifically, pro-inflammatory IFNγ^+^ CD4^+^ T cells were increased in KO ears **(Figure 2D),** suggesting an enhanced T_H_1 response consistent with an influx of macrophages (**Figure 2E).** Interestingly, T_H_17 cells were also expanded in KO mice skin samples **(Figure 2F)**, potentially promoting increased neutrophil recruitment **(Figure 2G)**.

The other tissue most affected by loss of *Tfrc* in *Foxp3*-YFP-Cre was the lungs. Pathology analyses revealed moderate to severe perivascular and peribronchiolar infiltrates, primarily composed of lymphocytes with fewer neutrophils, eosinophils, and plasma cells (**Figure 2H**). Indeed, flow cytometric quantification of immune cells confirmed CD4^+^ and CD8^+^ T cell infiltration in the lungs from KO mice (**Figure 2I**). To characterize the type of T cell inflammation in the lungs, transcription factors and cytokines were assessed from lung T cells. Similar to scurfy mice (37), KO mice had increased T_H_1 and T_H_2 cells (**Figure 2J and K**), although they also had increased T_H_17 cells in the lung (**Figure 2L**). Concomitantly, the frequency of lung-resident Treg cells was significantly reduced in KO mice, although still detectable at around 3% of the CD4^+^ T cell compartment (**Figure 2M**). Together, these data support a tissue-specific requirement of CD71, particularly in lung and skin Treg cells to maintain tolerance.

### Tfrc KO Treg cells are unstable

*Tfrc* KO Treg driven by the *YFP*-*Foxp3*-Cre were previously shown to have reduced markers of stability such as CTLA-4, ICOS, and PD-1 (11). Consistent with this, splenic Treg cells also showed a significant loss of stability markers Nrp-1 and Helios (**Figure 3A and B**). Conversely, KO Treg cells demonstrated increased CD39 expression, an ectoenzyme associated with Treg control of inflammatory autoimmune disorders (**Figure 3C**) (38). We then asked whether these unstable Treg cells express inflammatory transcription factors. Indeed, within the Foxp3^+^ T cell population, more Treg cells in KO mice co-expressed T-bet and RORγt than WT mice. (**Figure 3D and E**).

**Figure 3:**
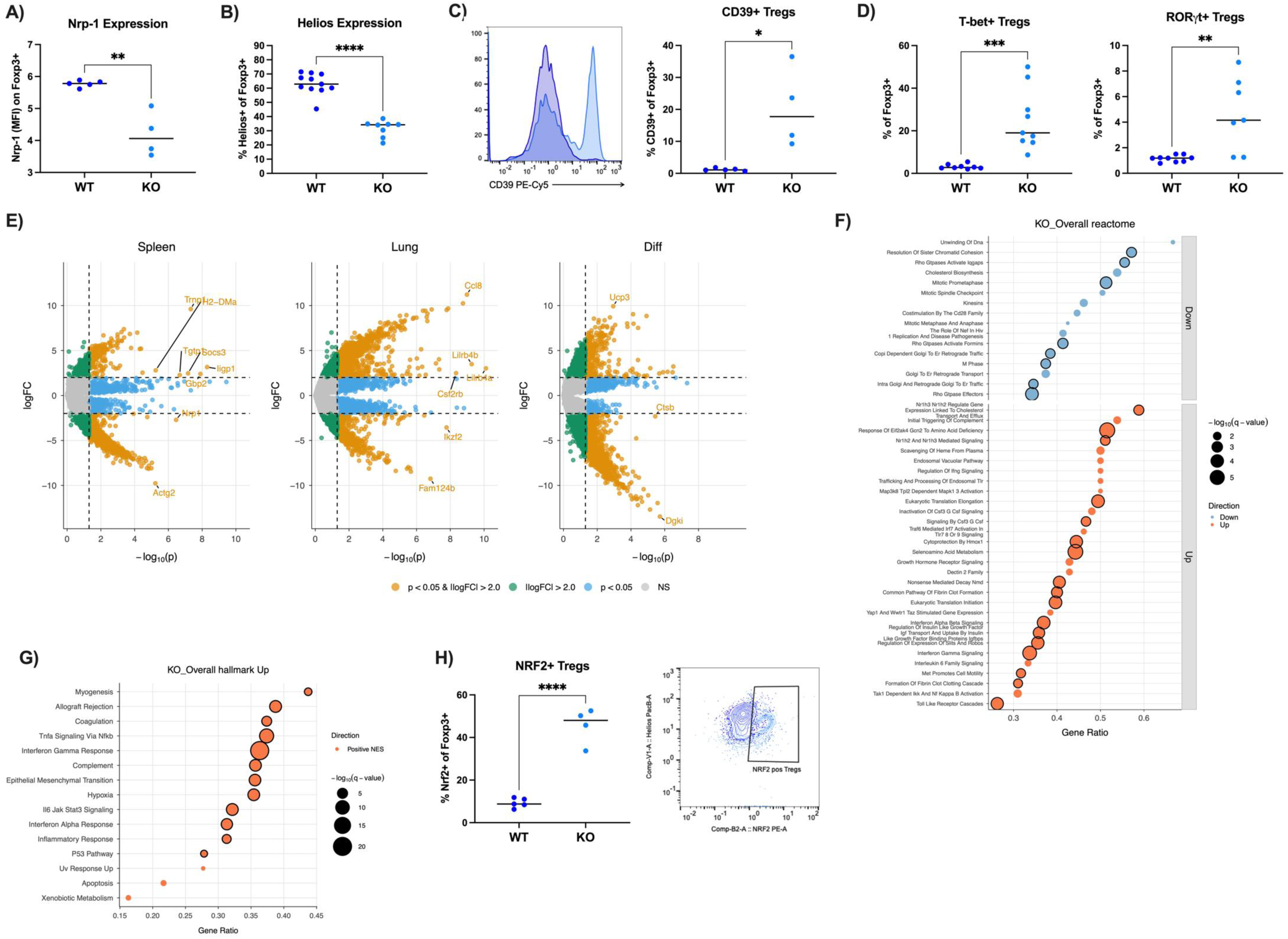
Loss of *Tfrc* in Tregs alters their stability. Spleens from Foxp3^YFP^-Cre, *Tfrc* floxed mice were processed into single cell suspensions for flow cytometry analysis of Treg markers (A) Nrp-1 (B) Helios and (C) CD39. D) Within the Foxp3+ population, cells positive for expression of T-bet or RORyt were quantified as percentages. E) RNA-sequencing analysis of Tregs sorted from spleen or lungs of Foxp3^YFP^-Cre, *Tfrc* floxed mice. Volcano plots for the three pre-specified contrasts fit with limma-voom. Left (Spleen): Spleen_KO − Spleen_WT. Middle (Lung): Lung_KO − Lung_WT. Positive logFC in the Spleen/Lung panels indicates higher expression in KO vs WT within that tissue. Right (Diff): genotype×tissue interaction (Spleen_KO − Spleen_WT) − (Lung_KO − Lung_WT). Positive logFC means the KO effect is stronger in Spleen than in Lung (negative = stronger in Lung). Points are genes; x-axis is −log10(p), y-axis is log2 fold-change (logFC). Vertical dashed line marks p = 0.05; horizontal dashed lines mark |logFC| = 2. NS = not significant. F) Dot plot showing Reactome pathways enriched for the KO_Overall contrast, defined as the pan-tissue KO effect (Lung_KO + Spleen_KO)/2 − (Lung_WT + Spleen_WT)/2. GSEA was run with fgsea (100,000 permutations) on all genes passing filterByExpr, ranked by the moderated t-statistic from Model 2. Points represent pathways; the x-axis is GeneRatio (size of the leading-edge/core-enrichment gene set divided by the pathway size); point size encodes −log10(q-value); color encodes direction (Up = positive NES, Down = negative NES). Black outlines indicate pathways with FDR q < 0.05. G) Hallmark pathways with positive normalized enrichment scores (NES) for KO_Overall (definition as above). GeneRatio on the x-axis, point size = −log10(q-value). Black outlines mark pathways passing FDR q < 0.05. H) Splenic Tregs as in A-D were analyzed for NRF2 expression (left). NRF2 expression was simultaneously analyzed with Helios expression within Foxp3+ cells (right).

To better characterize the functional defects in KO Treg cells in this model, YFP^+^ Treg cells were sorted from spleen and lungs of 3-week-old KO mice or their WT littermates. Bulk RNA-sequencing analyses of sorted Treg cells confirmed a loss of Helios (*Ikzf2*) and Nrp-1 (*Nrp1*) expression across tissue types (**Figure 3F**). Interestingly, the Reactome pathway analysis revealed many altered metabolic processes including cholesterol biosynthesis and transport as downregulated compared to WT Treg cells, and selenoamino acid metabolism and Gcn2 signaling to amino acid deficiency as upregulated (**Figure 3G**). Along with selenoamino acid metabolism which contributes to antioxidant defense, the Hallmark dataset genes associated with hypoxia were also significantly upregulated in KO Treg cells (**Figure 3H**). Together, these data indicate that oxidative stress could contribute to KO Treg cell destabilization. In support of this observation, KO Treg cells showed 4 to 5 times higher NRF2 expression levels than that of WT Treg cells (**Figure 3I**). Interestingly, Treg cells that were positive for Nrf2 expression also tended to be negative for Helios expression. Take together, these results show that *Tfrc* deletion compromises Treg cell stability, in part through increased oxidative stress and expression of proinflammatory transcription factors.

### Inducible Tfrc deletion drives tissue-specific inflammation

We next turned to a tamoxifen-inducible Foxp3-Cre-ER^T2^ model of *Tfrc* deletion in Treg cells to understand the role of CD71 expression in Treg cells in adults as opposed to during the development of tolerance early in life. Another advantage of this model is the opportunity to obtain larger cell numbers of purified WT or KO Treg cells. We first performed a full necropsy on mice 3 weeks after their first tamoxifen injection or solvent control (**Supplemental Figure 4**). A cohort of 3 WT and 3 KO males, and 1 WT and 1 KO female were evaluated. Mild to moderate extramedullary hematopoiesis (EMH) was observed in all KO mice and only one WT mouse within the spleen. Half of the KO mice showed mild white pulp atrophy. The tracheobronchial, mandibular, mesenteric, and axillary lymph nodes were examined. Minimal to mild sinus histiocytosis, defined as increased numbers of macrophages within the node sinuses, was observed in several KO animals. There was a mild increase in severity of sinus histiocytosis within the KO versus the WT mice, with some neutrophilic infiltrates and loss of lymphoid cells were noted in the KO mice. In the pancreas, mild multifocal mononuclear infiltrates were observed within the majority of the KO mice, which was not present in the WT mice.

Whereas pathology was noted in the spleen, LN and pancreas of KO mice, many tissues appeared within normal limits in both WT and KO mice including the brain, skeletal muscle (bicep), heart, thymus, bladder, bones (femur, sternum) and external ear. Only the pinna and shallow regions of the external ear were evaluated. Most surprising was the small and large intestines were microscopically within normal limits in the examined H&E-stained sections for all animals. This contrasts with previous studies which highlighted lesions within the gastrointestinal tract of these specific mice (10, 11).

The inducible KO model was far less severe than the constitutive *YFP*-Cre model, with no significant increase in spleen size (**Figure 4B**). Furthermore, in contrast to the constitutive model, mice treated with tamoxifen had no loss of Foxp3^+^ Treg cells within the CD4^+^ T cell compartment. Treg cell frequencies were instead increased in KO mice (**Figure 4C**). This was also true in the lungs and skin draining lymph nodes (dLN) (**Figure 4D and E**). Although lung pathology was present, there was no significant increase in CD4^+^ and CD8^+^ T cell frequencies in the lungs (**Figure 4F and G**). However, the CD8^+^ T cells present were more activated, suggesting a role in local inflammation (**Figure 4H and I**). Consistent with the constitutive KO model, Treg cells lost a significant amount of Helios expression and increased their co-expression of T-bet in the lung (**Figure 4J+K**). In summary, inducing Treg-specific *Tfrc* deletion in healthy adult mice causes Treg cell to adopt a proinflammatory and increased CD8^+^ T cell activity likely due to lower suppressive capacity.

**Figure 4.**
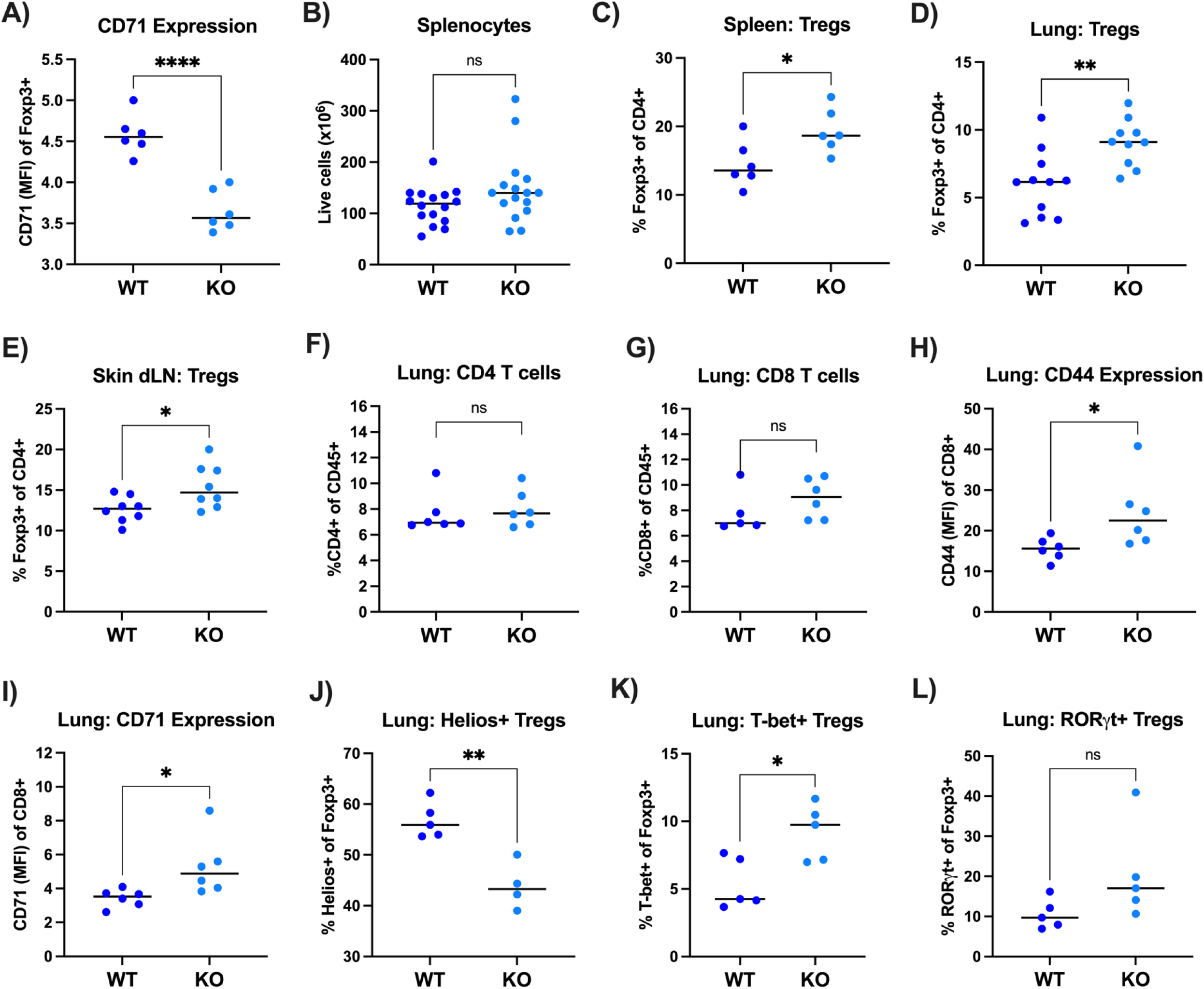
Loss of *Tfrc* in adults alters Treg frequencies and activation across tissues. 8 to 10 week-old Foxp3-EGFP-ER^T2^-Cre mice were treated with tamoxifen or corn oil injections for 2 weeks to induce deletion of *Tfrc*. Flow cytometric analysis of T cell populations in WT and KO mice (A–L). A) CD71 expression after 3 weeks of tamoxifen or control treatment in splenic Tregs from KO mice compared to WT. (B) Splenocyte counts performed with trypan blue stain. Frequency of Foxp3⁺ Tregs among CD4⁺ T cells in the spleen (C), lung (D), and skin-draining lymph nodes (dLN) (E). Total frequencies of CD4⁺ (F) and CD8⁺ (G) T cells in the lung. H) Activation markers CD44 (H) and CD71 (I) on lung CD8⁺ T cells. Percentages of lung Foxp3+ Tregs that were Helios⁺ (J), T-bet⁺ (K), and RORγt⁺ (L). All statistical tests are unpaired two-tailed Student’s t-test.

### Treg cells metabolically adapt to a loss of Tfrc

Treg cell metabolism is essential for differentiation, stability, and suppressive function (39–41). Therefore, we asked whether loss of *Tfrc* in these Treg cells could impact their metabolic programming and thereby cause a loss of these features. While glucose importer Glut-1 expression on Treg cells was previously shown to support Treg proliferation, it also decreased suppressive function and was associated with a loss of Helios expression (14). Consistent with a slight expansion of Tregs in ER^T2^ KO mice (**Figure 4C**), Glut-1 protein expression on splenic Treg cells was significantly increased (**Figure 5A**). We hypothesized that KO Treg cells adapt to a loss of iron by increasing glucose uptake and glycolysis. To test this, cells were extracted from skin dLN and metabolic dependencies determined by SCENITH flow cytometry (42). Cells were incubated with either DMSO, 2-DG to disrupt glycolysis, oligomycin A (oligo) to disrupt mitochondrial ATP synthase, or both 2-DG and oligo and the mean fluorescence intensity of anti-puromycin calculated from Foxp3^+^ Treg cells (**Figure 5B**). To our surprise, KO Treg cells were less dependent on glycolysis despite increased Glut-1 expression (**Figure 5C**). Furthermore, KO Treg cells had a reduced glycolytic capacity. Instead, fatty acid oxidation and amino acid oxidation (FAO & AAO) were significantly increased (**Figure 5C**), suggesting that KO Treg cells adapt to a loss of *Tfrc* by increasing their oxidation of fatty acids and amino acids.

**Figure 5:**
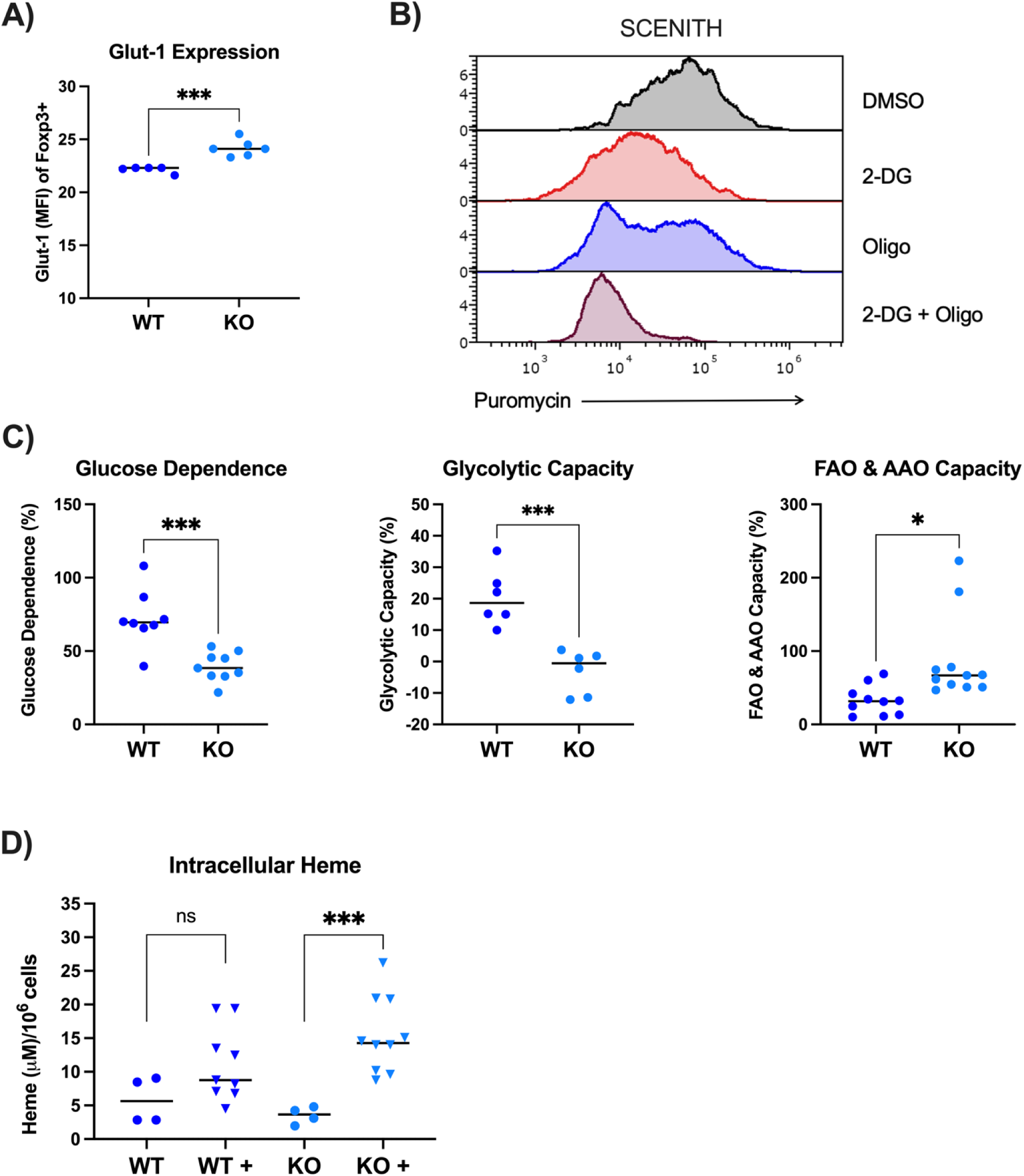
Metabolic Reprogramming of *Tfrc* KO Tregs. A) Glut-1 expression (MFI) of splenic Foxp3⁺ cells after tamoxifen or corn oil injections. B) Protein synthesis and thereby ATP levels were measured by SCENITH flow cytometry in skin dLN samples. After gating on Foxp3+ Tregs, mean fluorescent intensities (MFI) of anti-puromycin were quantified. MFI values from different metabolic inhibitor treatments were compared to the DMSO control to calculate metabolic dependencies and/or capacities. C) Metabolic dependency and capacity results from skin dLN Tregs. Significance by Student’s t-test. (D) Intracellular heme concentrations in purified WT and KO Treg cells with (+) or without incubation with exogenous heme. (A-C) unpaired two-tailed Student’s t-test, (D) One-way ANOVA with Sidak’s multiple comparisons test.

Because KO Treg cells in the ER^T2^ KO mice had increased frequencies (**Figure 4C-E**), we asked how these cells could adapt to proliferate without *Tfrc*-dependent iron uptake. Among significantly upregulated pathways from the Reactome database in the RNA-sequencing data were scavenging of heme from plasma and cytoprotection by heme oxidase 1 (Hmox1) (**Figure 3F**), suggesting increased heme uptake and metabolism. To test this hypothesis, Treg cells from WT and KO mice were purified and incubated with exogenous heme. After multiple washes, intracellular heme was measured, revealing increased heme concentrations after incubation (**Figure 5D**). Notably, KO Treg cells had a significant increase in heme whereas WT Treg cells did not. Together, these data suggest that KO Treg cells adapt metabolically to a loss of CD71-mediated iron uptake by increasing heme metabolism as an alternate iron source and increasing amino acid and fatty acid oxidation for ATP synthesis.

### Mice with Treg-specific Tfrc KO have exacerbated disease severity in atopic dermatitis model

Treg cells play an integral role in controlling allergic responses in addition to their role in preventing autoimmunity (43). We assessed CD71 expression and Tregs in a type 2 allergic disease model using the vitamin D_3_ analog Calcipotriol (MC903) to induce atopic dermatitis (AD) on the ears for 11 days (44). Indeed, the frequency of Tregs recruited to the ears of mice treated with MC903 was significantly higher compared to the vehicle control (**Figure 6A**) and animals treated with MC903 demonstrated enlargement of the draining parotid lymph nodes (pLN) (data not shown). Interestingly, CD71 expression on Tregs was higher in the pLN than in the ear tissue, and MC903 treatment resulted in more CD71 expression on pLN Tregs, suggesting an activated phenotype (**Figure 6B**). These Tregs also showed higher levels of labile, unbound iron, supporting a role for active iron metabolism in Tregs during AD disease (**Figure 6C**).

**Figure 6:**
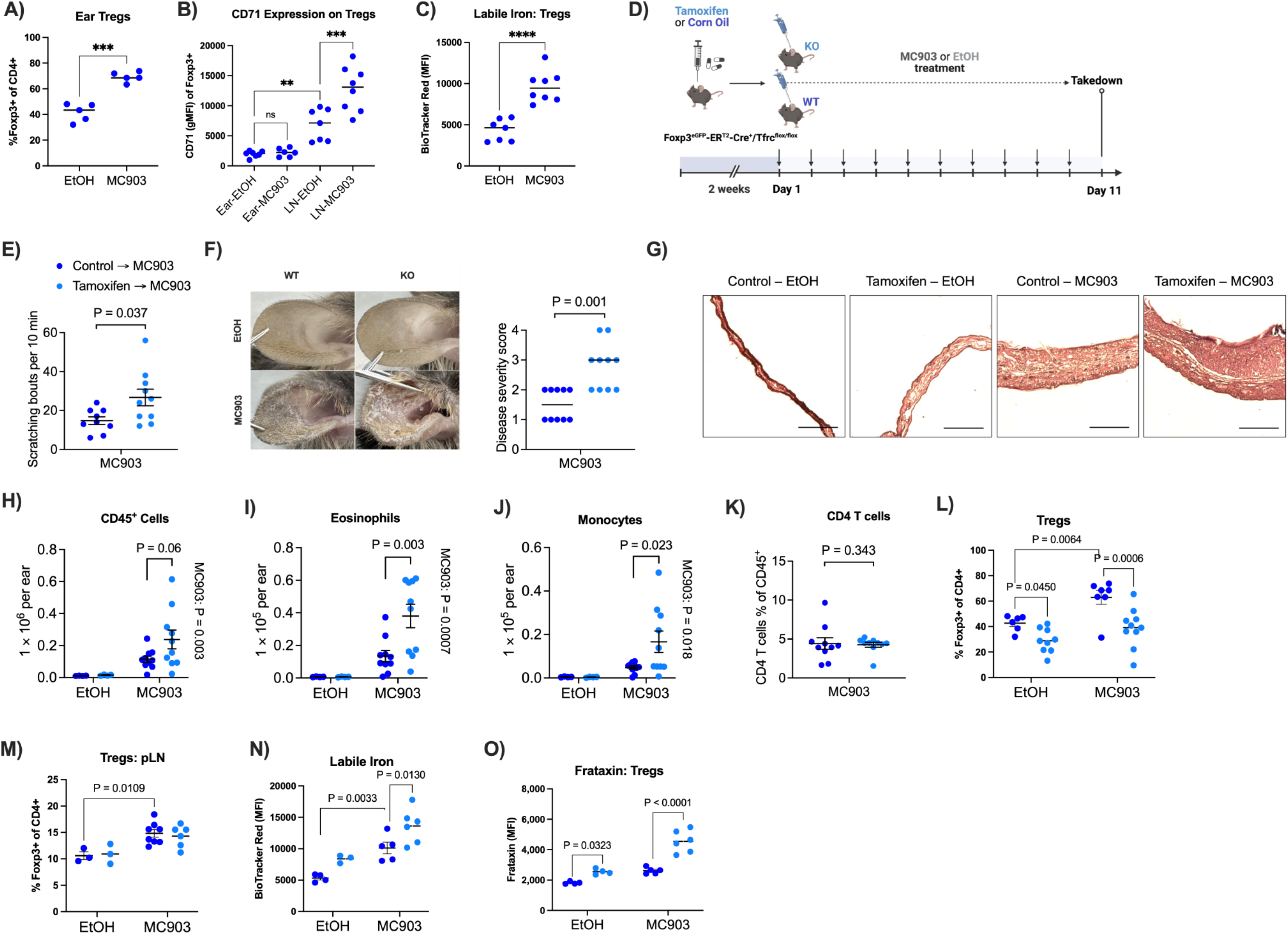
Tregs require *Tfrc* during atopic dermatitis. WT mice were treated with ethanol (EtOH) or MC903 for 10 days to induce atopic dermatitis (AD) on the ears. A) Frequency of Foxp3+ cells of CD45+CD4+ cells in the ears of WT mice. B) Flow cytometric quantification of geometric MFI (gMFI) of CD71 expression on Tregs from the ears or pLN of WT mice. C) BioTracker Labile Iron dye quantification of free iron in live GFP+ Tregs from the ears of WT mice. D) Foxp3-EGFP-ER^T2^-Cre mice were treated with tamoxifen or corn oil injections to induce deletion of *Tfrc* and subsequently treated with ethanol (EtOH) or MC903 for 10 days. E) On day 11, MC903 treated mice were blindly scored for scratching behavior for 10 minutes each. F) Representative images of ears from each treatment group (left). Disease severity scores were assessed by a blinded dermatologist based on Physician global assessment (PGA) score. G) Representative HE images of ears. Scale bar = 500 μm. H) Total CD45+ cells in the ears were counted by flow cytometry. I) Eosinophil counts defined as total live single cell CD45+ CD11b+ Ly6G- Siglec F+ cells. J) Monocyte counts defined as total live single cell CD45+ CD11b+ Ly6G- Siglec F- F480- CD11c-. K) CD4 T cell frequencies in MC903 treated mouse ears. L) Treg frequencies in the ears and (M) pLN. N) Free iron was quantified by flow cytometry in live GFP+ Tregs from the pLN. O) Frataxin expression of Foxp3+ Tregs was determined by flow cytometry in the pLN. A, C, E, G, H, M= Welch’s t test. B, H, I, J, L, M N, O= 2 way ANOVA.

Given this association with iron metabolism in Tregs during AD, we tested whether transferrin-bound iron is required for Tregs to respond to type 2 inflammation in this model using the tamoxifen ER^T2^ KO mice (**Figure 6D**). At day 10, itching behavior was assessed based on their scratching bouts and we observed increased itchiness in KO mice compared to WT (**Figure 6E**), indicating elevated disease severity. On day 11, we confirmed elevated disease severity in *Tfrc* KO mice by dermatologist-evaluated scoring of visual skin disease pathology (combining ear swelling, skin scaling and inflammation) and scaling (**Figure 6F**). Histological analysis of ears showed classical changes in epidermis (hyperkeratosis) and dermis (inflammatory infiltrates) associated with AD pathology in MC903-treated mice. Interestingly, we noticed more profound histological marks of dermal inflammation in MC903-treated KO compared to WT mice (**Figure 6G**), suggesting increased infiltration of myeloid cells.

To assess immune cell-mediated inflammation in ears, we quantified different immune populations of MC903-treated mice using flow cytometry. Overall, we found an increase in ear CD45^+^ cells in the *Tfrc* KO compared to WT mice (**Figure 6H**) consistent with a T_H_2-driven model of atopic dermatitis (45) mainly driven by elevated eosinophils and monocytes (**Figure 6I, J**). Despite similar frequencies of CD4 T cells (**Figure 6K**), Treg cells in the ears were decreased in *Tfrc* KO compared to WT mice (**Figure 6L**). Interestingly, the frequencies of Tregs in the pLNs were indistinguishable between KO and WT mice (**Figure 6M**), suggesting that recruitment and/or persistence in the ear tissue was specifically affected by a loss of *Tfrc*.

Tregs had increased levels of intracellular labile iron during AD challenge in the pLN (Figure 6C), so we hypothesized that *Tfrc* KO Tregs would have less intracellular iron which would hinder their suppressive capacity in AD. To our surprise, KO Tregs had higher levels of labile iron compared to WT Tregs in both the vehicle group and the MC903-treated mice (**Figure 6N**), suggesting that iron handling in KO Tregs is impaired at baseline. Consistent with high labile iron, expression of the mitochondrial iron chaperone protein frataxin was also increased in KO Tregs and was especially high after MC903 challenge (**Figure 6O**). Taken together, these results show that despite a less severe phenotype in adult mice, a loss of *Tfrc* disrupts intracellular iron handling in Tregs and ultimately impairs Treg function in type 2 inflammation in the skin.

## Discussion

Foxp3-expressing Tregs are central to the establishment of immune tolerance (46, 47) and play an integral role in shaping tissue repair, regeneration, and disease progression (48). Although tissue-resident Tregs share a similar identity across organs (49), they are also shaped by local microenvironmental cues such as nutrient availability and cell-cell interactions (40, 50). It is established that metabolism plays a central role from activation to suppressive function and stability of Treg cells (40, 51, 52) but less is known about how iron metabolism influences these qualities. In this study, we investigated the role of the transferrin receptor CD71 and iron metabolism in Treg cell stability and function with a focus on how cellular metabolism across different tissue niches could impact Treg biology.

We previously found that targeting CD71 on T cells with an antibody decreased T_H_1 and T_H_17 pathogenicity while promoting Treg Foxp3 expression, alleviating disease symptoms in a model of Systemic Lupus Erythematosus (SLE) (9). However, it was unclear whether the effect of CD71 blockade on effector T cells (T_eff_) indirectly contributed to the increased Treg frequencies in these mice, or it was a direct effect on Treg cells, or both. Moreover, the lethality of *Tfrc* deletion in mice makes it challenging to dissect Treg-specific effects from systemic inflammation feedback and signaling cascades in this severe phenotype. Thus, we compared tissue pathologies and Treg phenotypes in both the constitutive *Foxp3-*YFP-Cre model, and the inducible *Foxp3*-Cre-ER^T2^ model to investigate the importance of CD71 expression in different developmental stages, in both healthy and disease settings.

Previous studies established a critical role for CD71 in Tregs for establishing tolerance early in life, which was independent of thymic development and instead was microbiota-dependent (10, 11). In one study, Treg transfer experiments to *Rag1-/-* mice indicated an impaired ability of KO Tregs to prevent weight loss and intestinal inflammation compared to WT Tregs (11). However, here we report that using Tregs from a CD4-Cre *Tfrc* KO mouse model, KO Tregs were able to significantly suppress disease pathology in the same adoptive transfer/colitis model. We also did not observe significant gut inflammation in the *Foxp3* YFP Cre KO mouse model compared to WT littermates, suggesting that Tregs in the gut do not depend on CD71 expression for their function. Although these opposing outcomes could be influenced by the microbiome composition of different mouse colonies at different institutions, our results were consistent between two distinct institutions and mouse facilities.

Interestingly, *Rag1^-/-^* mice that received *Tfrc* KO Treg cells developed skin inflammation while the ones that received WT Treg cells did not, which suggested that Tregs may require *Tfrc* to maintain skin integrity or tolerance. This is supported by previously published single-cell RNA-seq data that we re-analyzed, showing high *Tfrc* expression in skin Treg cells compared to other tissues, pointing to their possible reliance on CD71 for their optimal function that tissue environment. Indeed, transferrin-bound iron availability in the skin is extremely scarce, with hair follicle progenitor cells relying on iron deliveries from iNKT cells (53). This would be in stark contrast to the gut microenvironment, where dietary iron absorption is taking place and iron should be available in multiple forms. Consistent with a requirement for CD71 expression in skin, KO mice in the *Foxp3* YFP Cre model also developed severe skin inflammation. Interestingly, this inflammatory phenotype was also observed in a previous study where Tregs were subject to iron overload and high CD71 expression (54), highlighting the careful tuning of iron levels required for optimal Treg function.

In our adult KO mice, however, we observed mostly lung pathology but no obvious signs of skin lesions. Nonetheless, we tested the function of KO Tregs in the skin of these adult mice using an inflammatory skin model of atopic dermatitis (AD) and found that mice with *Tfrc* KO Treg cells had exacerbated skin inflammation with higher eosinophil and monocyte infiltration levels and lower Treg cell numbers. The Treg cells lacking CD71 failed at containing effector T cells and thus could not resolve the inflammation or initiate tissue repair. In humans, skin Tregs express high levels of Arginase 2 (ARG2) which promotes OXPHOS and FAO while suppressing mTOR signaling (55). A loss of ARG2 expression in these Tregs is associated with a loss of suppressive function in psoriasis. Interestingly, ARG2 has also been implicated in the cellular regulation of immunosuppressive CD71^+^ erythroid cells in neonates (56). Therefore, it is tempting to hypothesize that in the skin, CD71-mediated iron metabolism is required for ARG2 activity which maintains an optimum balance of OXPHOS, mTOR and oxidative stress in Tregs.

Phenotyping of *Tfrc* KO Treg cells *ex vivo* confirmed that they have reduced stability markers such as Helios and increased expression of inflammatory transcription factors such as T-bet and RORγt. This correlated with the altered metabolic pathways such as decreased cholesterol biosynthesis and transport, which is crucial for the generation of Geranylgeranyl pyrophosphate (GGPP), a key metabolite that phosphorylates Stat5, which in turn is essential for Foxp3 maintenance in the gut (57). In general, disruption of cholesterol synthesis and lipid metabolism can significantly impair Treg cells making them more pro inflammatory (58). In addition, KO Treg cells showed signs of oxidative stress (high Nrf2), amino acid deprivation (GCN2 pathway enrichment) and low Helios, highlighting their instability. In a previous Treg study, excess intracellular iron caused by PDK1 deletion also caused increased oxidative stress and decreased Helios expression, ultimately destabilizing Tregs (54). Therefore, levels of intracellular iron whether too low or high may regulate Helios expression, or rather influence Treg stability via oxidative stress which is then indirectly marked by the presence or absence of Helios (59).

Higher oxidative stress could be a result of a dysregulated complex III of the respiratory chain that includes the Rieske iron-sulfur subunit, which is also the main source of ROS (60). Excessive ROS could be one of the causes of the instability, as loss of complex III leads to global DNA hypermethylation. Thus, hypermethylation could then negatively affect Foxp3 expression, suppressive capacity and thereby tolerance in the tissue. Indeed, iron metabolism was previously shown to regulate Treg stability through TET enzyme activity at the Foxp3 locus (39). Metabolically, *Tfrc* deletion rewired Treg cells to be more dependent on fatty acid oxidation and amino acid oxidation despite an increase in Glut-1 expression. To bypass the need for transferrin-bound iron uptake through CD71, *Tfrc* KO Treg cells were able to proliferate by increasing heme uptake as an alternate source of iron. Despite this increase in heme metabolism, it appears that this compensation was not sufficient to maintain full Treg stability and suppressive capacity, although the availability of heme for uptake would also differ depending on the tissue microenvironment.

Our findings provide evidence that depending on both CD71 levels and tissue location, Treg cells can be somewhat functional (colitis transfer model), completely dysfunctional (*Foxp3* YFP Cre model) or tolerate CD71 loss by maintaining suboptimal function (ER^T2^ inducible model). Although our mixed-use of models allows us to address CD71 dependencies of Treg cells during early development versus adult life, it has its challenges. *Tfrc* deletion efficiency via tamoxifen was not as robust, where Treg cells with CD71 expression are more likely to repopulate through selective pressure. Despite these limitations, we reveal a more complex picture of gut Treg biology and iron dependency than previous appreciated and implicate iron metabolism in Tregs in a new disease context of atopic dermatitis and type 2 immunity.

Despite lacking the receptor for transferrin-bound iron, KO Tregs were able to acquire some iron which appeared to occur at least in part through heme metabolism. Future research should examine questions such as how much iron and what form (labile, ferritin-bound, heme, etc.) is required for Treg function and available to Treg cells in different tissues. Indeed, we saw high intracellular labile iron in KO Tregs during allergic disease, which argues that iron levels alone do not predict cellular function.

Instead, the handling iron metabolism that dictates the balance of ferritin-bound versus labile iron in Tregs is critical for understanding their capacity. This is consistent with previous work which demonstrated that Tregs are especially dependent on higher ferritin expression to regulate iron stores compared to conventional T cells, required for *Foxp3* stability and suppressive function (39). This question becomes more complicated in the context of malnutrition or iron deficiency anemia, where a large portion of the global population is affected by dietary iron deficiency. On the opposite end of the spectrum, conditions of iron overload such as in primary and secondary hemochromatosis could impair Tregs through increased oxidative stress leading to cell death as in the PDK1 knockout cells (54). Overall, finding the right balance of iron uptake in different immune cell types at different developmental stages will be key to improve treatments targeting iron metabolism in these conditions.

## Abbreviations

Tfrc: Transferrin Receptor
T_H_: T helper
Treg: regulatory T cell
ROS: reactive oxidative species
CID: Combined immunodeficiency
WT: Wild Type
KO: Knockout
T_conv_: Conventional T cell
mLN: Mesenteric lymph nodes
pLN: parotid lymph nodes
T_FH_: T follicular helper
dLN: draining lymph nodes
FAO: Fatty acid oxidation
AAO: Amino acid oxidation
IP: Intraperitoneally
2-DG: 2-Deoxy-D-glucose
SCENITH: Single-cell energetic metabolism by profiling translation inhibition

