## Supplemental Figures for "Tissues Guide Dependence of Treg on the Transferrin Receptor"

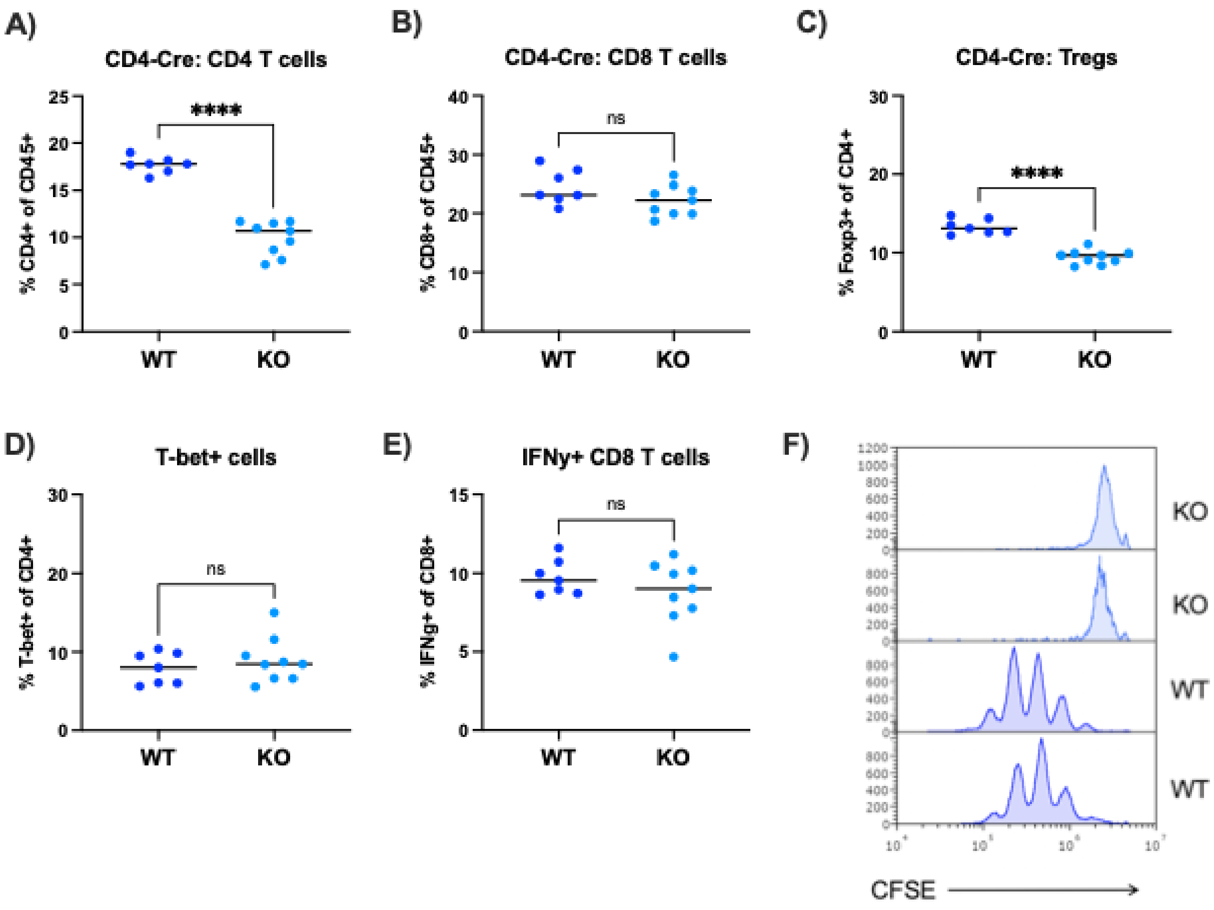


**Supplementary Figure 1. CD4-Cre *Tfrc*$^{fl/fl}$ KO mice do not exhibit T cell inflammation.**

Spleens were processed into single cell suspensions to analyze T cell populations by flow cytometry (A-E). Percentage of CD4+ (A) and CD8+ (B) T cells within total CD45+ cells. C) Percentage of Foxp3+ Tregs from Cre+ KO or Cre- WT mice. D) T-bet+ cell frequencies from WT and KO mice. E) Percentage of IFNy+ CD8 T cells after PMA and ionomycin stimulation. F) T cell proliferation of bulk CD4 T cells was assessed by CFSE dilution. Two representative WT and KO mice are shown after 3 days of activation. All statistical tests are unpaired two-tailed Student's t-test.

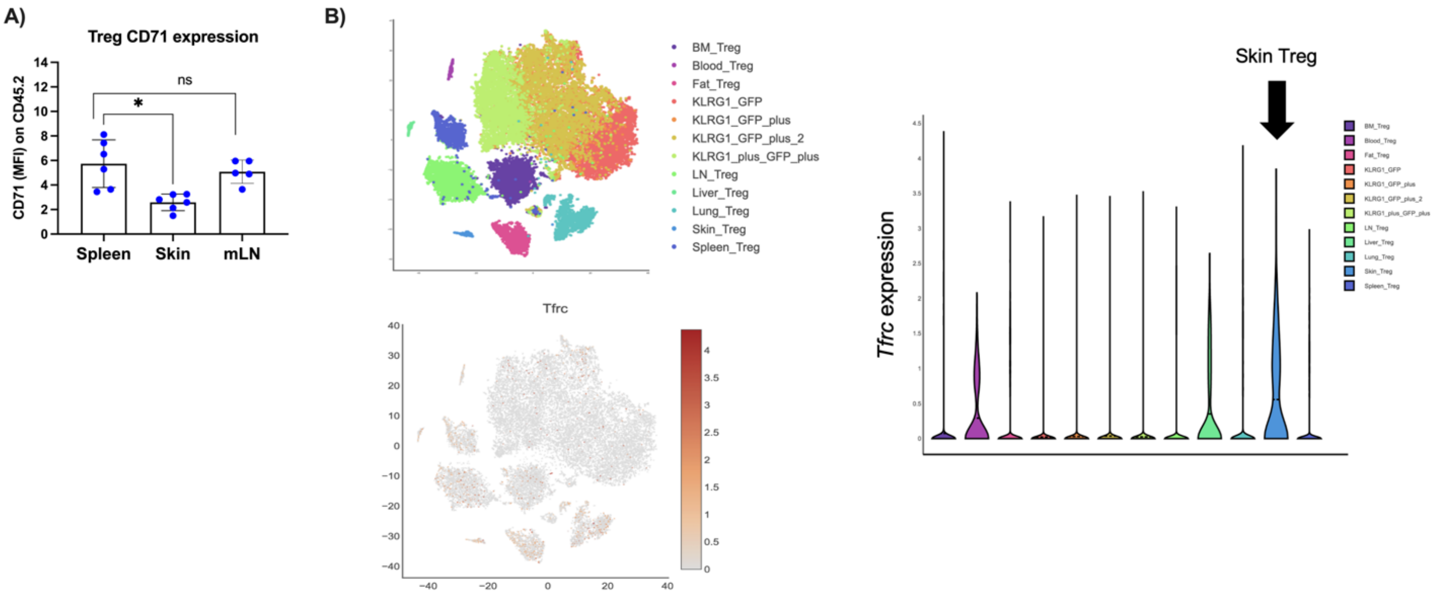


**Supplementary Figure 2. Tissue-specific expression of CD71 and *Tfrc* in Tregs.**

A) From the colitis model in Figure 1, CD71 mean fluorescent intensity (MFI) of WT transferred Tregs in spleen, skin, and mLN of recipient mice. B) Re-analysis of mouse Treg scRNA-seq dataset. Top left: t-SNE clustering of tissue specific Tregs, bone marrow (BM), blood, fat, lymph nodes (LN), liver, lung, skin, and spleen, etc. Bottom left: Feature plot showing expression of *Tfrc* across all Tregs. Right: Violin plot of *Tfrc* expression of tissue-specific Treg clusters with elevated expression in skin Tregs.

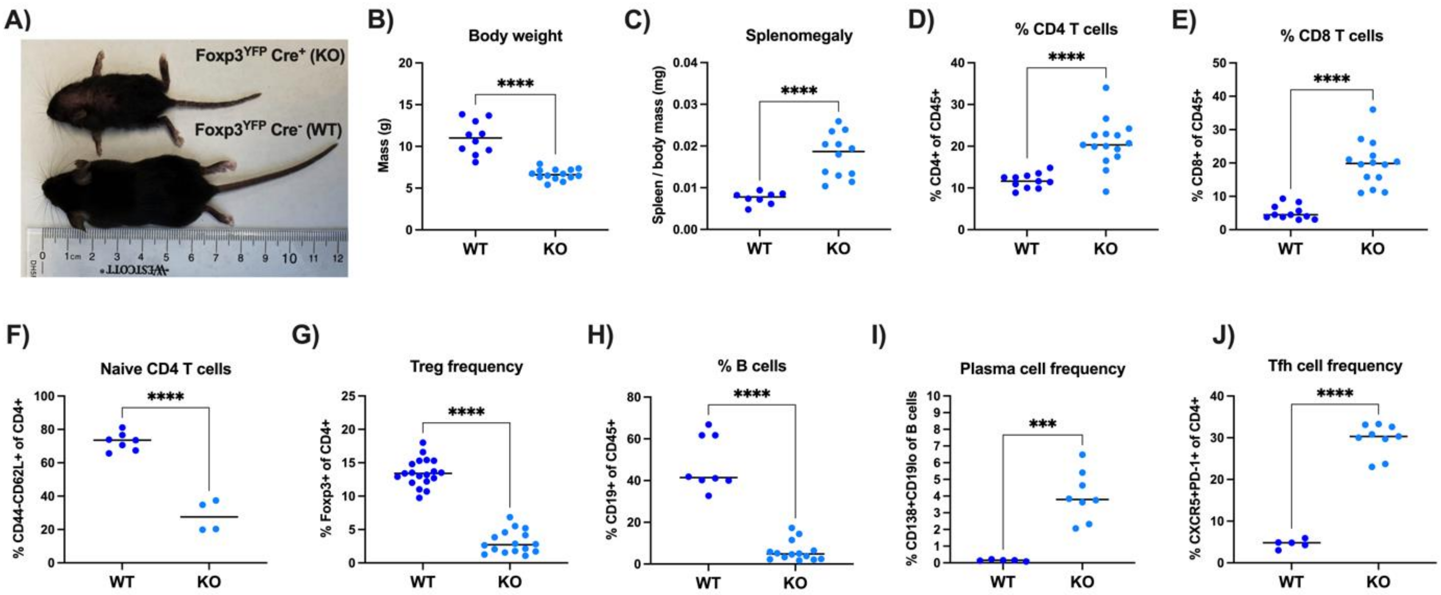


**Supplementary Figure 3. Foxp3$^{YFP}$-Cre; *Tfrc* floxed KO mice develop fatal autoimmune-like inflammation by 3 weeks of age.**

A) WT and KO littermates were observed for autoimmune phenotypes at 2.7 to 3 weeks of age. B) KO pups failed to thrive and did not develop healthy body weights. C) Enlarged spleens in KO mice. D) The frequencies of CD4 T cells (D) and CD8 T cells (E) in the spleens were determined by flow cytometry. F) Naive CD4 T cells and Treg frequencies (G) within the CD4 T cell population. H) Splenic B cell percentages and plasma cell frequencies (I) within the B cell population. (K) T follicular helper cell (Tfh) frequencies within CD4 T cell population. All statistical tests are unpaired two-tailed Student's t-test.

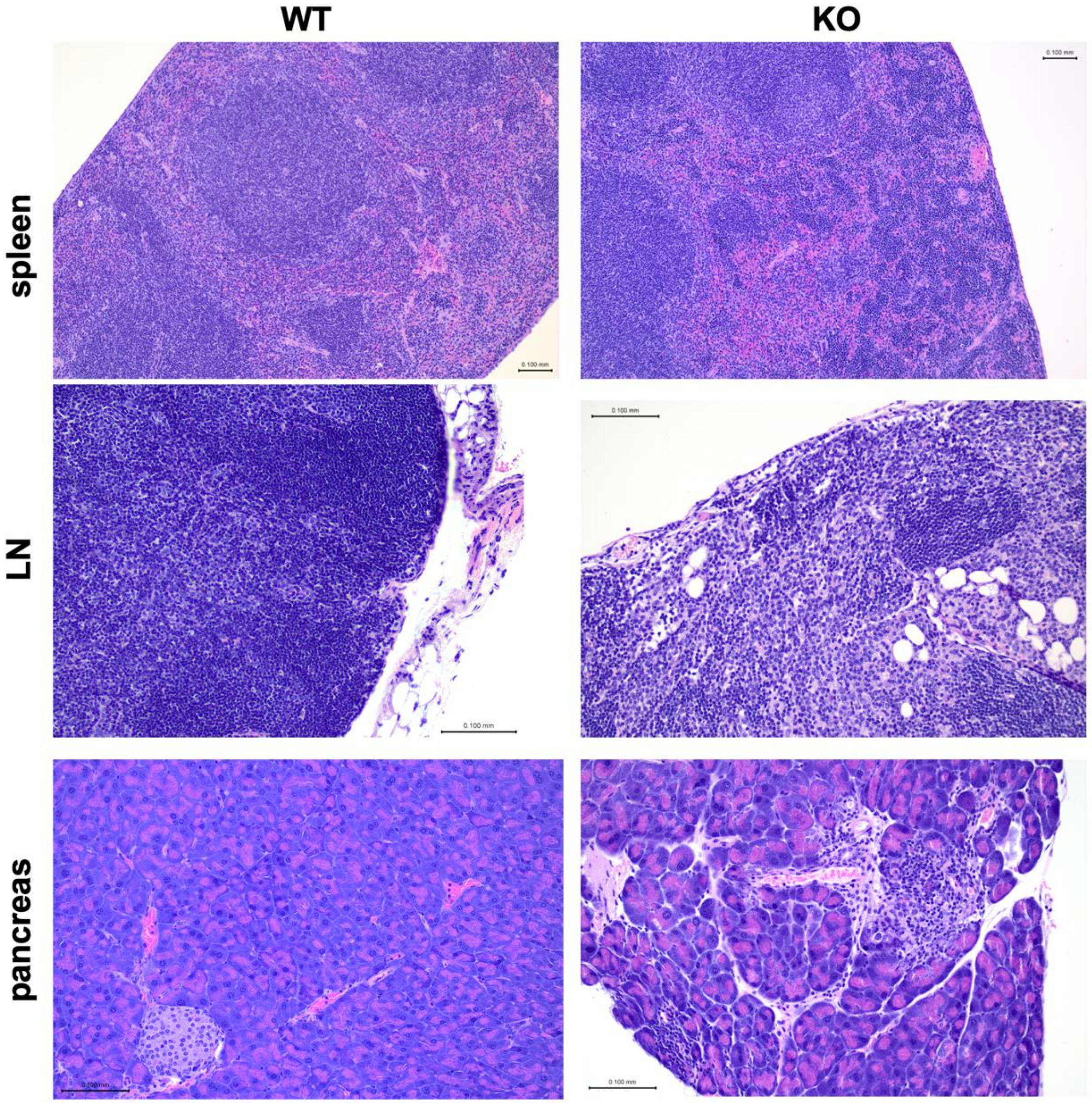


**Supplementary Figure 4. Histologic examination of tissue-specific inflammation in adult KO mice.**

8 to 10 week-old Foxp3-EGFP-ERT2-Cre mice were treated with tamoxifen or corn oil injections for 3 weeks to induce deletion of *Tfrc*. Representative images are shown of hematoxylin and eosin (HE)-stained spleens, lymph nodes (LN), and pancreas of 1 female WT and 1 female KO mouse.
